# Expanding the chemical diversity of RNA by transcriptional incorporation of amino acid- and glycosyl-modified nucleotides

**DOI:** 10.64898/2026.04.22.720138

**Authors:** Julián Valero, Kevin Neis, Laia Civit, Søren Fjelstrup, Maria Gockert, Jørgen Kjems

## Abstract

With the increasing interest in RNA-based therapies, there is a pressing need to incorporate new chemistries into more complex RNA molecules. These modifications can protect RNA from degradation, improve its pharmacokinetics, and enhance its targeting properties. Here we describe the enzymatic synthesis of chemically modified RNA derivatives using a mutant T7 RNA polymerase to incorporate 23 different base modifications alongside stabilizing ribose modifications, such as 2′-fluoro and 2′-deoxy groups. To investigate the impact on transcription efficiency and fidelity, we employed a pool of 38 template sequences and analyzed the transcripts by next-generation sequencing of the cDNA. Results demonstrated that all modifications were successfully incorporated into RNA, with transcription efficiency influenced by three main factors: type of modification, base modified, and the sequence context. Misincorporation levels during transcription and reverse transcription into cDNA were generally low (<1%) but included noticeable exceptions for some nucleobase-modification combinations. As a robust proof-of-concept we demonstrated the selection of Histidine-U modified aptamer, relying on multiple rounds of transcription and amplification, binding Influenza hemagglutinin protein with low nanomolar K_D_. We anticipate that this work will significantly contribute to the design and production of chemically modified RNAs with novel functionalities, advancing applications in biomedicine and synthetic biology.

**Graphical Abstract:** 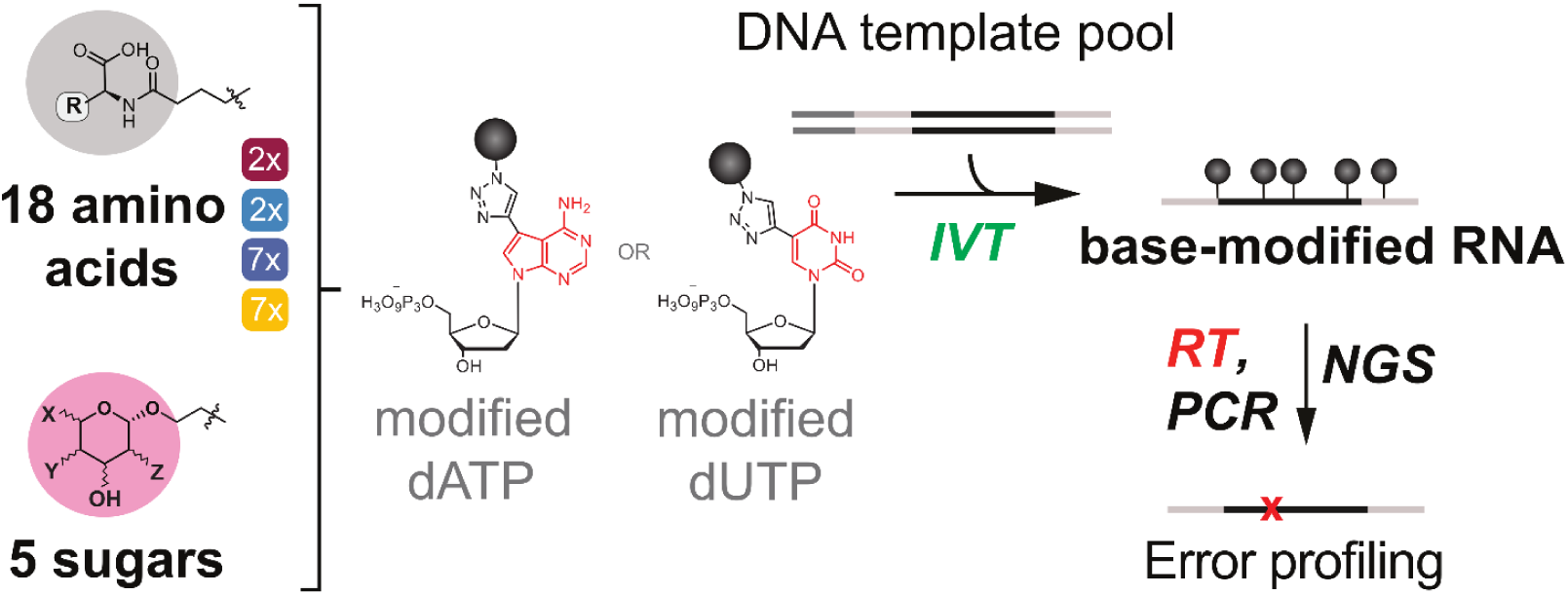

## INTRODUCTION

The functional and structural versatility of RNA in nearly all corners of life has led to considerable traction of this molecule for diagnostic and therapeutic applications, including mRNA, antisense oligonucleotides, CRISPR guide RNAs, RNA interference-based therapeutics, and RNA aptamers [1–5]. These diverse approaches share a common trait: the use of chemical modification to protect the RNA from degradation, enhance its pharmacokinetic profile and convey tissue/cell-specific targeting properties with reduced immunogenicity [5–7]. All RNA-based drugs currently on the market employ chemical modifications, targeting the phosphate backbone, the ribose sugar, or the nucleobase [7].

Backbone modifications focus largely on phosphorothioates, characterized by the substitution of the non-bridging oxygen with sulfur [8]. Ribose modifications mainly involve the 2’ position, either by replacement of the hydroxy group with other groups such as 2’-fluoro (F) or 2’-O-methyl, or by bridging the 2’ and 4’ ribose positions to form locked nucleic acids [7]. Nucleobase modifications can be divided into naturally occurring analogues and synthetic functionalization. Natural analogues, such as N1-methylpseudouridine, pseudouridine or 5-methylcytidine are commonly used to enhance stability and alleviate immune stimulation [9–12]. Synthetic modifications, especially on pyrimidines, are used to introduce positionally defined functional groups into RNAs, including fluorophores, affinity tags and reactive groups [13–21]. These functionalized RNAs offer diverse applications ranging from RNA imaging and real-time single-molecule monitoring of biological processes to crosslinking of protein complexes. In addition, base modifications can notably enhance the targeting properties and binding affinity, conferring attractive advantages for theranostic development. For example, amino acids groups in DNA have allowed to drastically extend the range of accessible protein targets for aptamers, advancing biomarker discovery [22,23]. Next to amino acids, glycosyl modifications have been applied to improve the targeting properties [24], as exemplified by the broad use of N-acetylgalactosamine (GalNAc) siRNA conjugates for targeting the Asialoglycoprotein receptor 1 (ASGPR1) [25]. Generally, glycan-lectin interactions are dependent on multivalent and spatial organization of the glycosyl groups making them interesting candidates for programmable nucleic acid platforms [26,27].

To date, most studies featuring enzymatic or post-synthesis base-modification focus on DNA, whereas available methods and accessible chemical diversity for RNAs are more limited [19,22,28–30]. Chemically modified RNAs can be produced synthetically via solid-phase synthesis, but this approach suffers from high costs, incompatibility with certain modifications and, importantly, remains practically limited to molecules shorter than 40-60 nucleotides (nts) [31]. Also, for applications where large pools of RNAs with different sequences need to be synthesised from mixed templates (e.g. for in vitro selection of RNA), chemical synthesis is not an option. For these applications, enzymatic synthesis offers a scalable alternative. Besides enzymatic ligation [32] and primer-extension reaction with engineered DNA polymerases [20,33,34], RNA transcription using recombinant polymerases (mainly T7, T3, SP6 and mutants thereof) is widely employed [35–38]. The efficiency of these enzyme-driven in vitro transcriptions (IVTs) of chemically modified RNA depends strongly on the nature of the chemical modification. While IVT-mediated integration of simple sugar or base modifications has been investigated and optimized, more complex and unconventional modifications have been less studied [39].

Here we report a method for transcriptional synthesis of RNAs with efficient and accurate incorporation of nucleotides containing a large variety of base-coupled amino acid- and sugar-functional groups, alongside with ribose modifications, such as 2′-deoxy and 2′-fluoro modifications known to enhance stability against degradation [40–45]. Comprehensive comparison of 18 different amino acid and 5 sugar modifications on either deoxy-adenine or deoxy-thymine demonstrated base- and modification-dependent effects for Y639F mutant T7 RNA polymerase (T7-RNP) mediated incorporation efficiency. Furthermore, detailed NGS based error profiling/ fidelity assessment using a DNA template pool of 38 different 80-82-nts sequences, demonstrated high fidelity transcription and reverse transcription reactions, with relatively low-rate substitutions, especially transversions, as the main error type, and homopolymeric regions as error hotspots. Closer assessment of error sources revealed that reverse transcription is significantly more error-prone than in vitro transcription. Finally, an exemplary proof-of concept application highlighted the robustness of the presented method, yielding successful in vitro selection of Histidine-U modified aptamers exhibiting high affinity towards Influenza hemagglutinin. The presented work expands the chemical diversity of RNA, holding great promise for applications in biomedicine, RNA technology and synthetic biology.

## MATERIALS AND METHODS

### General remarks

All synthetic oligonucleotides (primers, splint, ssDNA templates, and KK_RNA_BioAdapt) were synthesized by Integrated DNA Technologies (IDT). Copper(II) sulfate (CuSO₄), magnesium chloride (MgCl₂), HEPES, acetonitrile, sodium acetate, dithiothreitol (DTT), spermidine-HCl, bovine serum albumin (BSA), triethylammonium acetate (TEAA) buffer, ascorbic acid, sugar-azides, amino acids, dimethyl sulfoxide (DMSO), and PEG 8000 were purchased from Sigma-Aldrich. BTTAA, canonical NTPs, and pCp-Biotin were obtained from Jena Bioscience. Alkyne-modified dNTPs (ethynyl-dUTP and ethynyl-dATP) were supplied by baseclick GmbH. Nuclease-free water, canonical dNTPs, SuperScript III reverse transcriptase, and salmon sperm DNA were purchased from Invitrogen. Azidobutyric acid NHS ester was obtained from Lumiprobe, and 2′-F-dCTP was purchased from TriLink BioTechnologies. Inorganic pyrophosphatase (IPP), DNase I, RiboLock RNase inhibitor, Tween-20, Phusion High-Fidelity DNA Polymerase, *BamH*I, GeneJET PCR Purification Kits, Klenow Fragment (exo-), and T4 RNA ligase were purchased from Thermo Fisher Scientific. D-PBS (10×) was obtained from Gibco. RNA 5′ pyrophosphohydrolase (RppH), NEBuffer 2, and T4 RNA Ligase 2 were purchased from New England Biolabs. The Y639F mutant T7 RNA polymerase was obtained from Aptamist Aps.

### Synthesis of chemically modified dNTPs

#### Azide-functionalization of amino acids

Azide functionalities were introduced to the amino acids by reacting the respective amino acid (3.6 µmol, 3 equiv, in H_2_O) with azidobutyric acid NHS ester (1.2 µmol, 1 equiv, in DMSO) in HEPES buffer (50 mM, pH 8.2, 5% DMSO) at a final volume of 200 µL. The reaction mixture was incubated at room temperature for 5 h or at 4 °C overnight.

#### CuAAC coupling of azido molecules to alkyne-modified dNTPs

Sugar-azide derivatives and azidefunctionalized amino acids were conjugated to the alkyne-modified dNTPs via copper-catalyzed azide-alkyne cycloaddition (CuAAC). First, a 250 µL CuAAC catalyst buffer was prepared by incubating CuSO_4_ (1.25 µmol, 1 equiv, in H_2_O) with BTTAA (2.50 µmol, 2 equiv relative to CuSO_4_, in DMSO) for 5 min at room temperature. A freshly prepared ascorbic acid solution (40 µmol, 32 equiv, in H_2_O) was then added to the buffer mixture.

For the sugar modifications, sugar-azides (2.55 µmol, 3 equiv, in DMSO) were mixed with either 5-Ethynyl-dUTP or 7-deaza-7-Ethynyl-dATP (0.85 µmol, 1 equiv, in H_2_O) and the pre-mixed CuAAC catalyst buffer in HEPES buffer (50 mM, pH 7.4, 5% DMSO) at a final volume of 500 µL. For the amino acid modifications, the pre-mixed crude azide-functionalized amino acids (1.2 µmol azide groups, 3 equiv) were mixed with either 5-Ethynyl-dUTP or 7-deaza-7-Ethynyl-dATP (0.4 µmol, 1 equiv, in H_2_O) and the pre-mixed CuAAC catalyst buffer in HEPES buffer (50 mM, pH 7.4) at a final volume of 500 µL.

The final reaction mixture was incubated under light exclusion at 30 °C for 2 h. The products were purified using reverse-phase high-performance liquid chromatography (RP-HPLC). A Kinetex C18 column (Phenomenex) on a 1260 Infinity II HPLC system (Agilent) and an acetonitrile/H_2_O/50 mM TEAA mobile phase were used for purification. Product-containing fractions were monitored at 287 nm, lyophilized, reconstituted in nuclease-free water, and their concentrations were determined by measuring the absorbance at 287 nm.

### In vitro transcription, RNA PAGE purification and quantification

#### In vitro transcription protocol

The transcriptions were performed in transcription buffer (80 mM HEPES, pH 7.5, 25 mM MgCl_2_, 2 mM spermidine-HCl) supplemented with BSA (0.05 mg/mL), DTT (12.5 mM), inorganic pyrophosphatase (0.005 U/µL), and DMSO (10% v/v, or 15-20% v/v for hydrophobic modifications). All reactions contained 2’F-dCTP and GTP (2.5 mM each). Control reactions contained canonical ATP and UTP (2.5 mM each), while reactions containing modified dATP or dUTP (1.25 mM) replaced the respective canonical NTP. PCR-amplified DNA templates (0.7 µM) and Y639F mutant T7 RNA polymerase (1 U/µL) were added last. The transcription mixture was incubated for 5 h at 37°C.

#### RNA PAGE purification and quantification

Transcription reactions were incubated with DNase I (0.07 U/µL) for 30 min at 37 °C. Afterwards, samples were purified on an 8% denaturing polyacrylamide gel and recovered via passive elution in sodium acetate (0.3 M, pH 5.2) and subsequent ethanol precipitation. Relative quantification of the transcription was done by measuring the absorbance of the purified samples at 260 nm using the formula:

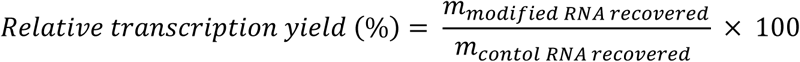

### Biotin-RNA adapter annealing and cycling reverse transcription (RT) on immobilized RNA

Relevant DNA sequences are listed in table S3.

#### RNA preparation

Target RNA (reference sequence 19, see Supplementary Information) was transcribed and purified as described above. Transcription reactions used ATP, 2′F-dCTP, GTP, and either canonical UTP or the modified analogues His-dUTP, Leu-dUTP, or Phe-dUTP. To generate the 5′-monophosphates required for downstream ligation, purified RNA transcripts were incubated with RNA 5′ Pyrophosphohydrolase (2.5 U/µg RNA) in 1× NEBuffer 2 for 30 min at 37 °C and subsequently purified using the RNA Clean & Concentrator-25 Kit (Zymo Research).

#### Splinted ligation of the biotinylated UMI adapter

For adapter ligation, the 5′-monophosphorylated RNA, a biotinylated RNA adapter containing a unique molecular identifier (KK_RNA_BioAdapt), and a complementary DNA splint were mixed in equimolar ratio (1 µM each). The mixture was heated to 65 °C for 3 min and rapidly cooled on ice for 3 min to facilitate annealing. Ligation was performed using T4 RNA Ligase 2 (0.5 U/µL) in 1× T4 RNA Ligase 2 buffer supplemented with PEG8000 (10% w/v), MgCl₂ (8 mM), and RiboLock RNase inhibitor (1 U/µL). The reaction was incubated for 2 h at 25 °C, followed by heat inactivation at for 5 min at 80 °C.

#### Cycling reverse transcription

Ligated RNA was immobilized on Dynabeads™ MyOne™ Streptavidin C1 (Invitrogen) according to the manufacturer’s protocol, using 1× D-PBS supplemented with 0.01% Tween-20 for all binding and washing steps. Following the removal of unbound RNA, the beads were resuspended in nuclease-free water containing 0.01% Tween-20. Reverse transcription primer (KKrv_short, 5 µM) and dNTPs (0.5 mM each) were added to the bead-bound RNA, incubated for 5 min at 65 °C, and rapidly cooled down on ice to facilitate annealing. Subsequently, First-Strand Buffer (1×), DTT (5 mM), RiboLock RNase inhibitor (1 U/µL), and SuperScript III reverse transcriptase (10 U/µL) were added. Reverse transcription was performed for 45 min at 50 °C. The synthesized cDNA was eluted by heating the mixture at 85 °C for 4 min, and the supernatant containing the eluted cDNA was immediately separated from the bead-bound RNA using a pre-chilled magnetic rack. To generate multiple independent cDNA copies for downstream UMI-family analysis, the RNA-immobilized beads were resuspended in nuclease-free water containing 0.01% Tween-20, fresh reverse primer (KKrv_short), and dNTPs and subjected to subsequent rounds of annealing and reverse transcription as described above. This cycling RT procedure was repeated for a total of three RT rounds.

### SELEX protocol

DNA sequences relevant for the SELEX protocol are listed in table S5.

#### Initial RNA SELEX library generation

Selection of histidine-modified RNA aptamers was performed using a library (termed N40-mU) consisting of a 40-nt random region flanked by two constant regions lacking uridines. dsDNA transcription templates were generated via Klenow extension. Specifically, the forward primer (N40_mU_fw) was annealed to the ssDNA library (N40_mU_ssDNA) at a 1.2-fold molar excess. Annealing was performed by heating the mixture to 95 °C for 5 min, cooling to 50 °C for 5 min, and then holding at 37 °C with a ramp rate of 0.1 °C/s. Extension was carried out using the Klenow Fragment (exo-) at 37 °C for 1 h. The resulting dsDNA library was purified by 6% native polyacrylamide gel electrophoresis (PAGE), followed by passive elution in sodium acetate (0.3 M, pH 5.2) and ethanol precipitation. The initial RNA library was generated and PAGE-purified following the transcription protocol described above, except that the dsDNA template was used at a final concentration of 1.2 µM, and canonical UTP was replaced with His-dUTP (0.625 mM).

#### Target immobilization and initial selection cycle

Selection was performed against a His-tagged H1 A/England/195/09 (H1) recombinant protein immobilized on Dynabeads™ His-Tag Isolation and Pulldown magnetic beads (Invitrogen). In the first selection cycle, 7.5 µg of H1 protein was incubated with magnetic beads (1:1.25) ratio for 15 min at room temperature with continuous rotation in PBS (pH 7.4) supplemented with Tween-20 (0.01%). The beads were washed three times with washing buffer (WB: PBS, pH 7.4, 1 mM MgCl_2_), then resuspended in selection buffer (SB: WB supplemented with 0.1 mg/mL salmon sperm DNA and 0.1 mg/mL BSA). An equivalent amount of magnetic beads without protein was prepared for counter-selection. Prior to selection, 1.7 nmol of the initial RNA library was refolded in WB using the following thermal protocol: 90 °C for 2 min, 65 °C for 5 min, 37 °C for 5 min, and cooling to room temperature. The refolded library was first incubated with the uncoated counter-selection beads for 45 min at room temperature with continuous rotation. Unbound sequences (supernatant) were then transferred to the H1-protein-coated beads and incubated under identical conditions. The beads were washed three times with WB (0.5 mL per wash, 1 min each). Bound RNA was reverse transcribed directly on the beads using the reverse primer (N40_mU_rv) and SuperScript III reverse transcriptase. The resulting cDNA was PCR amplified using Phusion High-fidelity DNA Polymerase and the specific primers (N40_mU_fw and N40_mU_rv), and the dsDNA was purified using the GeneJET PCR purification kit.

#### Iterative selection cycles

For subsequent selection cycles, the His-modified RNA library was transcribed as described above and purified using the RNA Clean & Concentrator-25 kit. The input RNA library amount was reduced to and maintained at 200 pmol for all subsequent rounds. The amount of immobilized H1 target protein was gradually decreased to increase selection pressure: 4 µg in cycle 2, 1.5 µg in cycles 3 and 4, and 0.75 µg in cycles 5 and 6. Starting from cycle 2, a more stringent counter-selection step was introduced to eliminate His-tag binders, utilizing 4 µg of an unrelated His-tagged nanobody immobilized on beads. The counter-selection procedure from cycle 2 onward was as follows: the RNA library was incubated with the nanobody-coated beads for 25 min at room temperature with rotation. The supernatant was subsequently incubated with uncoated beads for 10 min under identical conditions, followed by final incubation of the supernatant with the target H1-protein-coated beads. Incubation time with the target beads was reduced to 10 min for later selection cycles (cycles 4-6), and washing steps were progressively extended to increase stringency.

### NGS library preparation and sequencing

Relevant DNA sequences are listed in table S3.

Three distinct next-generation sequencing (NGS) libraries were prepared to evaluate: (i) Combined IVT and RT fidelity and sequence bias for each modification, (ii) IVT-templated errors via UMI-family analysis, and (iii) aptamer enrichment across SELEX rounds. The sample preparation for each library is described here:

#### (i) Combined IVT and RT fidelity and sequence bias for each modification

Samples were prepared for sequencing following an adapted protocol [46]. RNA was reverse transcribed using SuperScript III reverse transcriptase and a specific reverse primer (KKrv). Next, dsDNA was generated via PCR using Phusion High-Fidelity DNA Polymerase and specific forward (KKfw) and reverse (KKrv) primers. The resulting amplicons were digested with *BamH*I, purified using a GeneJET PCR Purification Kit, and subsequently PCR-amplified using primers containing sequencing adapters and unique index sequences. The final PCR products were purified using the GeneJET PCR Purification Kit and pooled in equimolar amounts.

#### (ii) IVT-templated errors via UMI-family analysis

For each sample, cDNA from the three cycling RT reactions was PCR amplified using specific forward (KKfw_adapter) and reverse (KKrv_short) primers with Phusion High-Fidelity DNA Polymerase. The resulting dsDNA was extended in a second PCR using forward (KKfw_ex-tension) and reverse (KKrv_extension) extension primers. Each library was subsequently indexed and adapter-extended via PCR using the Nextera XT Index Kit v2 Set A (Illumina). The final PCR products were purified using the GeneJET PCR Purification Kit and pooled in equimolar amounts.

#### (iii) Aptamer enrichment across SELEX rounds

DNA libraries from selection rounds 2-6, as well as the initial histidine-modified RNA library (post-RT-PCR), were extended using specific extension primers (Overhang_fw and Overhang_rv). A subsequent PCR was performed to incorporate unique index sequences and Illumina sequencing adapters using the Nextera XT Index Kit v2 Set A. The final PCR products were purified using the GeneJET PCR Purification Kit and pooled in equimolar amounts.

#### Final Library Processing and Sequencing

For all three experiments, the pooled libraries were size-selected on a Pippin Prep instrument using a 3% agarose, 100–250 bp cassette (Sage Science) to remove excess adapter primers. Library size distributions and purity were confirmed by agarose gel electrophoresis, and final concentrations were determined using a Qubit fluorometer (Invitrogen). Final sequencing libraries were prepared according to the manufacturer’s guidelines for the iSeq 100 System (Illumina). Sequencing was performed on the iSeq 100 System.

### NGS data analysis

#### Preprocessing and quality control

For the overall error analysis (i), initial preprocessing was performed using an in-house software pipeline [46]. For the IVT-templated error and aptamer enrichment analyses (ii and iii), preprocessing of single-read R1 reads was performed on the Galaxy server (https://usegalaxy.eu/). Quality control of FASTQ files was assessed using FastQC (v0.11.9, http://www.bioinformatics.babraham.ac.uk/pro-jects/fastqc/). Adapter and constant regions were trimmed using Cutadapt (v4.0) [47] with default settings, and low-quality reads were filtered using fastp (v0.23.2) [48] with a minimum Phred score of 30. Identical sequences were subsequently collapsed to determine raw read counts. All downstream analyses were performed using custom Python scripts (v3.10.9), which are available upon reasonable request. A summary of each analysis workflow is provided below.

#### (i) Combined IVT and RT fidelity and sequence bias for each modification

NGS reads were assigned to one of the 38 reference sequences (“Seeds”, Table S2) based on minimal global edit distance (Levenshtein distance, LD), using the *rapidfuzz* package (v3.13.0). Reads with a LD ≤ 5 to a single unique seed were retained. This filtering yielded between 106,287 and 371,003 reads (median = 208,258) per sample. Replicate agreement was assessed by comparing seed abundance profiles across both replicates using Pearson correlation coefficients and the slope of replicate-to-replicate proportional abundance. Replicates were pooled for all down-stream analyses. Deviations in relative sequence abundances within the modified sample pools, compared to their respective ATP and UTP control samples, were quantified using the Jensen-Shannon divergence [49]. Per-base composition profiles were computed for each sample relative to the respective control.

Reads were aligned to their assigned seeds using a global Needleman-Wunsch alignment. For each unique sequence, the read count, presence of substitutions, insertions, and deletions, as well as the substitution pattern, substitution positions, and flanking bases, were recorded. Reads were initially grouped by the non-exclusive presence of any of the three error types to quantify overall error presence. Substitution, insertion and deletion rates were calculated after filtering out reads containing two or more different error types. Error rates were defined as the number of error events divided by the total number of analyzed bases and reported as errors per 1000 bases. Substitution patterns (reference base → observed base) and local sequence context (`NNYNN` around the substituted base) were determined. All rates were weighted by the read counts of each unique sequence.

#### (ii) IVT-templated errors via UMI-family analysis

Preprocessed reads were parsed to extract the UMI, constant regions, and variable region. Passing reads were required to meet the following criteria:

1. Presence of the 5’ anchor sequence (`ATCC`) with a LD ≤ 1, followed by a 9-nt UMI.
2. Correct alignment to the internal constant region (GCCGAGTGCAGCGGGGCCACCAACGACATT) with an alignment score ≥ 25 (via Biopython *PairwiseAligner*) and a correct starting position.
3. Variable region length ≥ 20 nt.
4. Variable region matching reference sequence 19 with an overall LD ≤ 10, and an LD ≤ 2 within the first 5 nucleotides.

The parsing yielded between 119,135 and 177,733 passing reads (median = 157,810) per sample. Sequences sharing the same UMI across RT replicates were grouped into families of size 1, 2, or 3, corresponding to their occurrence in one, two, or three independent RT samples. For duplicated UMIs within a single RT sample, only sequences representing ≥85% of the reads for that specific UMI were retained. Within each family, substitutions (position and identity), insertions and deletions relative to reference sequence 19 were determined using *rapidfuzz*. Variable regions containing insertions or deletions were excluded from subsequent analysis. Substitutions were classified as follows:

- *Size 1 family substitution:* UMI family detected in only one RT reaction.
- *One-sided substitution:* UMI family detected in 2 or 3 RT reactions, but the substitution was present in only one of those replicates.
- *Transcription-based substitution:* UMI family of size 2 or 3 exhibiting identical substitution patterns across at least 2 independent RT reactions.

#### (iii) Aptamer enrichment across SELEX rounds

Preprocessed sequences, yielding between 129,623 and 307,981 reads (median = 161,591) per sample, were combined, clustered, sorted by read count, and filtered to retain only sequences with > 10 reads. The LD between sequences was calculated using *jellyfish* (v0.9.0), starting from the most abundant sequences. Sequences with an LD < 5 were merged by summing their read counts and retaining the most abundant representative sequence. Enrichment was calculated as the fraction of a sequence’s read count over the total pool read count. Base distributions across selection rounds were computed by summing read-weighted nucleotide counts at each position.

### Biolayer interferometry (BLI) binding assays

#### 3′-Biotinylation of RNA aptamers

Selected aptamer sequences (2 µM) were first biotinylated at the 3’-end using T4 RNA ligase (0.3 U/µL) and pCp-Biotin (6 µM). The ligation reaction was performed in 1× T4 RNA ligase buffer supplemented with ATP (1.5 mM), PEG 8000 (15 % w/v), inorganic pyrophosphatase (0.001 U/µL), and incubated overnight at 4 °C with vigorous shaking. Biotinylated RNA was subsequently purified using the RNA Clean & Concentrator-25 kit, and final RNA concentrations were determined by UV-Vis spectroscopy.

#### BLI measurements

Binding and binding affinity of the selected aptamer sequences to the H1 A/England/195/09 hemagglutinin trimer were assessed using the Octet®Red96 system (ForteBio). All measurements were performed in 96-well black, flat-bottom plates (Greiner) with orbital shaking set to 1,000 rpm. Refolded, biotinylated RNA aptamer were immobilized on Octet® Streptavidin (SA) Biosensors (Sartorius). For initial screening, the loaded biosensors were dipped into a 150 nM solution of H1 trimer in BLI buffer (1× D-PBS supplemented with 1 mM MgCl_2_, 0.1 mg/mL BSA and 0.01% Tween-20). Specificity was evaluated using 150 nM of a related hemagglutinin protein (H5 A/whooper swan/Mongolia/244/2005) and 300 nM of the His-tagged nanobody previously used for counter-selection. To assess the role of the histidine modifications in aptamer binding, control RNA aptamers transcribed with alternative modified nucleotides (2′F-dUTP or Leu-dUTP replacing His-dUTP) were biotinylated and tested as under identical conditions. For kinetic affinity measurements, loaded biosensors were dipped into a twofold dilution series of H1 trimer starting from a maximum concentration of 150 nM. All BLI experiments followed a standardized protocol: baseline acquisition in BLI buffer, an association phase in the protein solution (150 s for screening, 400 s for kinetic curves), and a dissociation phase in BLI buffer (100 s for screening, 1200 s for kinetic curves). A fresh biosensor was used for each protein concentration. Sensorgrams were baseline-aligned, and the data were analyzed using the instrument’s built-in software and EVILFIT [50,51].

## RESULTS

### Synthesis and enzymatic incorporation of amino acid– and glycosyl-modified dNTPs

A diverse set of base-modified deoxynucleotide triphosphates (dNTPs) bearing amino acid and sugar moieties were synthesized. Eighteen different amino acids (all except proline and lysine) and five different sugars were included presenting chemical groups of varying size, polarity, and charge. Azide groups were first introduced to amino acids by coupling the α-amino groups to a bifunctional azidobutyric acid *N*-hydroxysuccinimide (NHS) ester. These azido-functionalized amino acids, along with azide-containing sugars, were subsequently conjugated to either C5-ethynyl-dUTP or C7-ethynyl-7-deaza-dATP nucleotides through copper(I)-catalyzed azide-alkyne cycloaddition (CuAAC) (Fig. 1A). The positioning of the modifications on the Hoogsteen edge of the nucleobase is expected not to interfere with Watson-Crick base pairing. All modified dNTPs bearing amino acid and sugar moieties were successfully synthesized and characterized by LC-MS after HPLC purification (Table S1).

**Figure 1:**
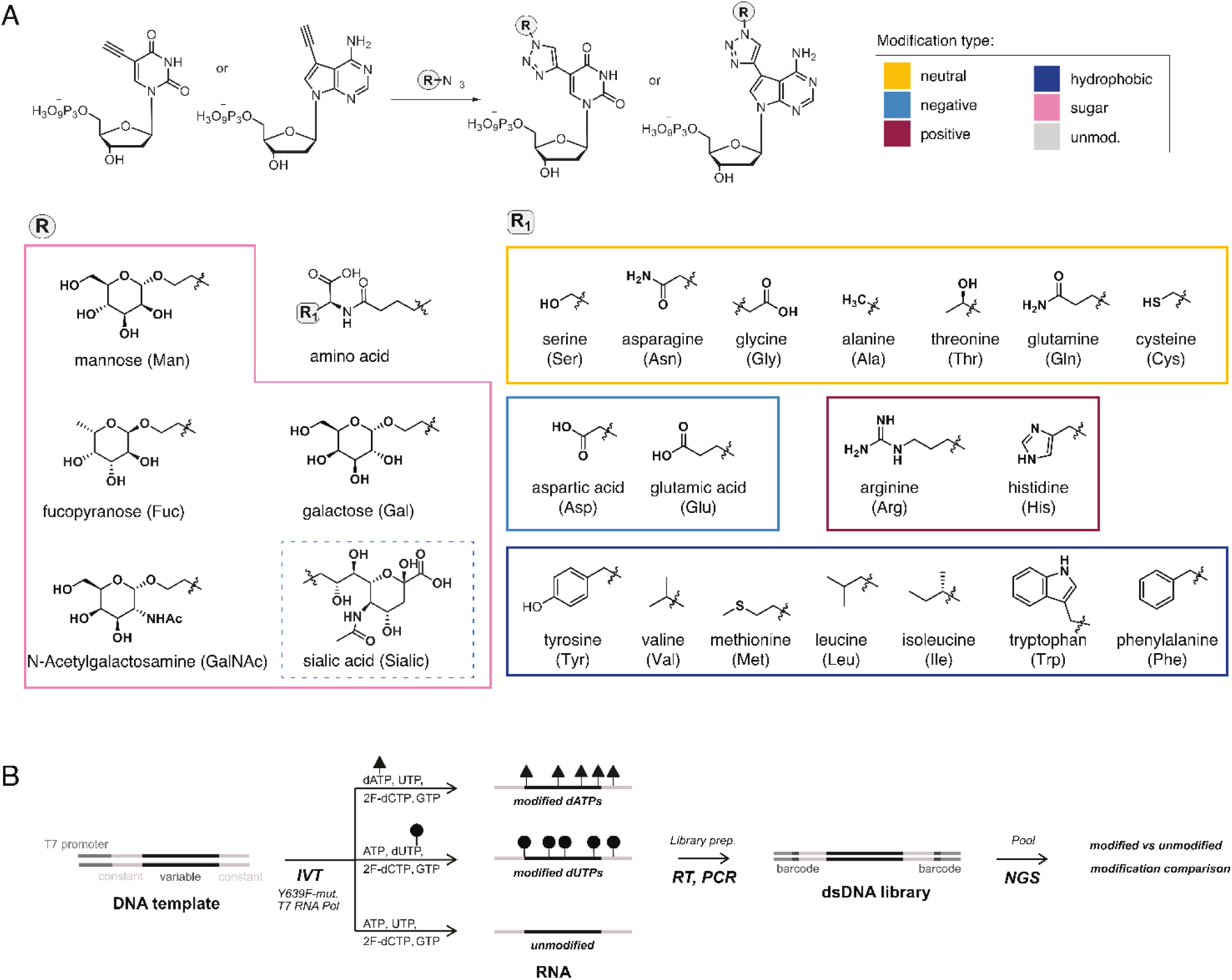
**A)** Click reactions forming the base-modified nucleotides and chemical structures of all modifications (R). Sugar modifications (pink box) were coupled via an azide functionality to alkyne functionalized deoxynucleotide triphosphates (dNTPs) using Copper(I)-catalyzed azide–alkyne cycloaddition (CuAAC). Amino acid residues (R1) were first coupled at the α-amino group to an azide-containing linker by NHS chemistry and then reacted with the dNTPs via CuAAC. Panel colour represents modification type, with neutral amino acid modifications in yellow, negatively charged in light-blue, positively charged in dark-red and hydrophobic in dark-blue. **B)** Experimental design to study transcription efficiency and accuracy of individual test sequences: A Y639F-mutant T7 RNA polymerase is employed for IVT of a pooled DNA template library (n=38) using nucleobase-modified dATP or nucleobase-modified dUTP or unmodified ATP/UTP, together with 2′F-CTP and GTP. Resulting RNA pools are reverse transcribed using SuperScript III, PCR-amplified using primers containing modification and sample-specific barcodes, and sequenced on an Illumina iSeq 100 platform. For protocol and sequence details see Supplementary Materials. Modified adenosines are indicated by round points, modified uridines by triangles.

Next, we assessed the incorporation efficiency of this modification panel into RNA using the Y639F mutant T7 RNAP, which was selected for its established tolerance towards both 2’-ribose (e.g., 2’F and 2’-deoxy) and base modifications [36]. The 46 modified dNTPs were evaluated as substrates for transcribing a pool of 38 defined DNA templates, each comprising a variable region of 35–37 nts, flanked by two constant regions of 18 nt and 27 nt at the 5′- and 3′ end, respectively (Table S2). The resulting RNA transcripts (80-82 nt in total) were designed to maximize sequence diversity, with variable adenosine and uridine content (20-33%) and structural motifs such as A-rich or U-rich regions and homopolymer stretches. Base-modified transcripts were generated by fully substituting either canonical ATP or UTP with the corresponding modified dATP and dUTP, respectively. Additionally, all reactions included 2′F-dCTP, commonly used for enhanced nuclease stability, and GTP. Modifications of guanosine were avoided due to the requirement for guanosine at the +1 position during T7 transcription initiation and control transcriptions were performed without base-modified nucleotides.

All modified nucleotides except threonine-dUTP yielded the desired transcription products. Quantification of full-length RNA yields after PAGE purification, relative to controls without base modifications (Fig. 2A), revealed that incorporation efficiency by the Y639F T7 RNAP strongly depended on both the nucleobase and the nature of the modification. In general, C5-modified dUTPs yielded on average 13% more full-length RNA relative to C7-modified dATPs. dUTPs modified with neutral, negatively charged, and hydrophobic amino acid modifications were generally incorporated more efficiently whereas modified dATPs favored sugar moieties, most notably mannose, fucose (high-yielding for both nucleobases), and *N*-acetylgalactosamine (GalNAc) (Fig. S1A, B). Furthermore, RNA yields tended to decrease with increasing amino acid steric bulk or hydrophobicity (Fig. S1C-F) and were further influenced by the amino acid charge: positively charged residues (arginine, histidine) were on average incorporated more efficiently than negatively charged ones (aspartic acid, glutamic acid), particularly when coupled to dATP.

**Figure 2:**
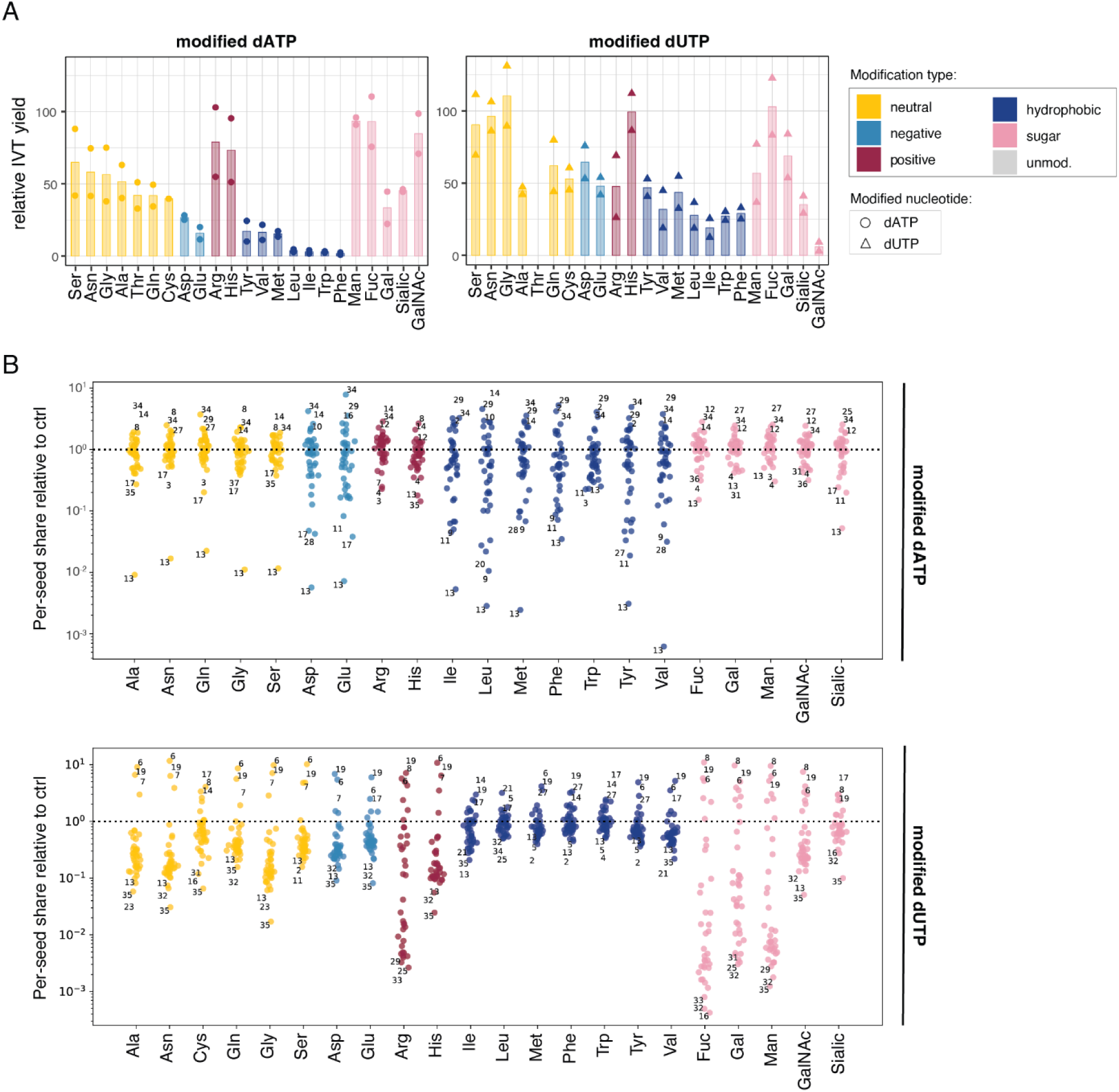
**A)** Relative in vitro transcription (IVT) yields from the pooled DNA templates using modified dATPs (left panel) or modified dUTPs (right panel). Yields were measured after PAGE purification and normalized to reactions containing unmodified rATP and rUTP, respectively. Bar plots show average values of two individual replicates except for Cys-dUTP (n=1). **B)** Per-seed enrichment for all assigned reads relative to control: For each sample the fraction of each sequence of the 38 sequences relative to the controls is plotted and the top and bottom three sequence IDs are shown. Logarithmic scale used. For Arg-, Fuc-, and Gal-dATP sequence 35 was not found. Dotted line = 1.

### Fidelity of transcription and reverse transcription of modified nucleotides

Having demonstrated the variable incorporation efficiencies among the tested modifications, we next evaluated their impact on sequence pool integrity and fidelity of transcription and reverse transcription. Because current direct RNA sequencing technologies are compatible with only a limited set of naturally occurring non-canonical base modifications [52,53], the fidelity of the combined transcription-reverse-transcription process was analyzed by evaluating sequence errors in the cDNAs generated by SuperScript III (SSIII) (Fig. 1B). This workflow enabled the identification of specific sequence motifs associated with reduced fidelity and the quantification of combined transcription–reverse transcription error rates.

Comparison of sequence pool composition across two replicates using Pearson correlation coefficients confirmed reproducibility (Fig. S1G). The impact of specific base modifications on library composition was examined at the individual sequence level (Fig. 2B) and at the global modification level using a sequence divergence metric (Fig. S1H). Overall, dUTP modifications induced a pronounced downward shift in abundance across most sequences within the pool, while a small subset became enriched, resulting in high divergence values (up to 0.6; Fig. S1H, right panel). In contrast, modified dATP samples maintained a more balanced read distribution, with relative abundances remaining closer to the controls and substantially lower global divergence (< 0.2; Fig. S1H, left panel). Notably, this pattern is not mirrored in the transcription yields (Fig. 2A). For modified dATPs, sequence abundance was most strongly affected with decreased transcription yield, most pronouncedly for negatively charged and hydrophobic amino acid residues. In contrast, modified dUTPs did not follow this trend. In spite of the low yield seen for hydrophobic modifications, the library composition remained similar to the control, while efficiently incorporated modifications, such as fucose, glycine and histidine, resulted in substantial variation compared to the control.

A more detailed sequence analysis of the copied cDNA highlights sequence-dependent effects on abundance shifts. Several of the test sequences consistently showed reduced or increased abundance across multiple modification types and, in some cases, independent of the modified nucleobase. To identify the underlying features driving these biases, we analyzed the three most and three least abundant cDNAs among the 38 test sequences across all modifications (labeled in Fig. 2B, Table S4). This analysis revealed that low-abundance sequences did not directly correlate with the total number of modified nucleotides in the transcript. Instead, a clear enrichment of homopolymer stretches comprising three or four consecutive modified A or T nucleobases was observed. This trend was particularly pronounced for modified uridines, where low-abundance sequences frequently displayed extended T-homopolymers. Notably, all low-abundance sequences began with one or two Ts, which, together with the two terminal Ts of the 5′ constant region, formed homopolymer tracts of three to four consecutive Ts. In contrast, high abundance sequences generally lacked extended homopolymer stretches, even when overall A/T content was comparable. This suggests that the occurrence of three or more modified uridines has a general negative impact on yields.

### Transversion events are most prevalent errors and generally associated with homopolymer stretches

The combined transcription-reverse transcription fidelity was assessed for each modification by categorizing the number of sequencing reads containing substitutions, insertions, or deletions relative to the reference sequence (Fig. 3A). Overall, high fidelity was maintained across most modified RNAs, with at least 82% of full-length reads containing no errors. A notable exception was observed for glycosyl-modified dATPs, which displayed a broad fidelity range, from 83% error-free reads for fucose to low 28% for sialic acid.

**Figure 3:**
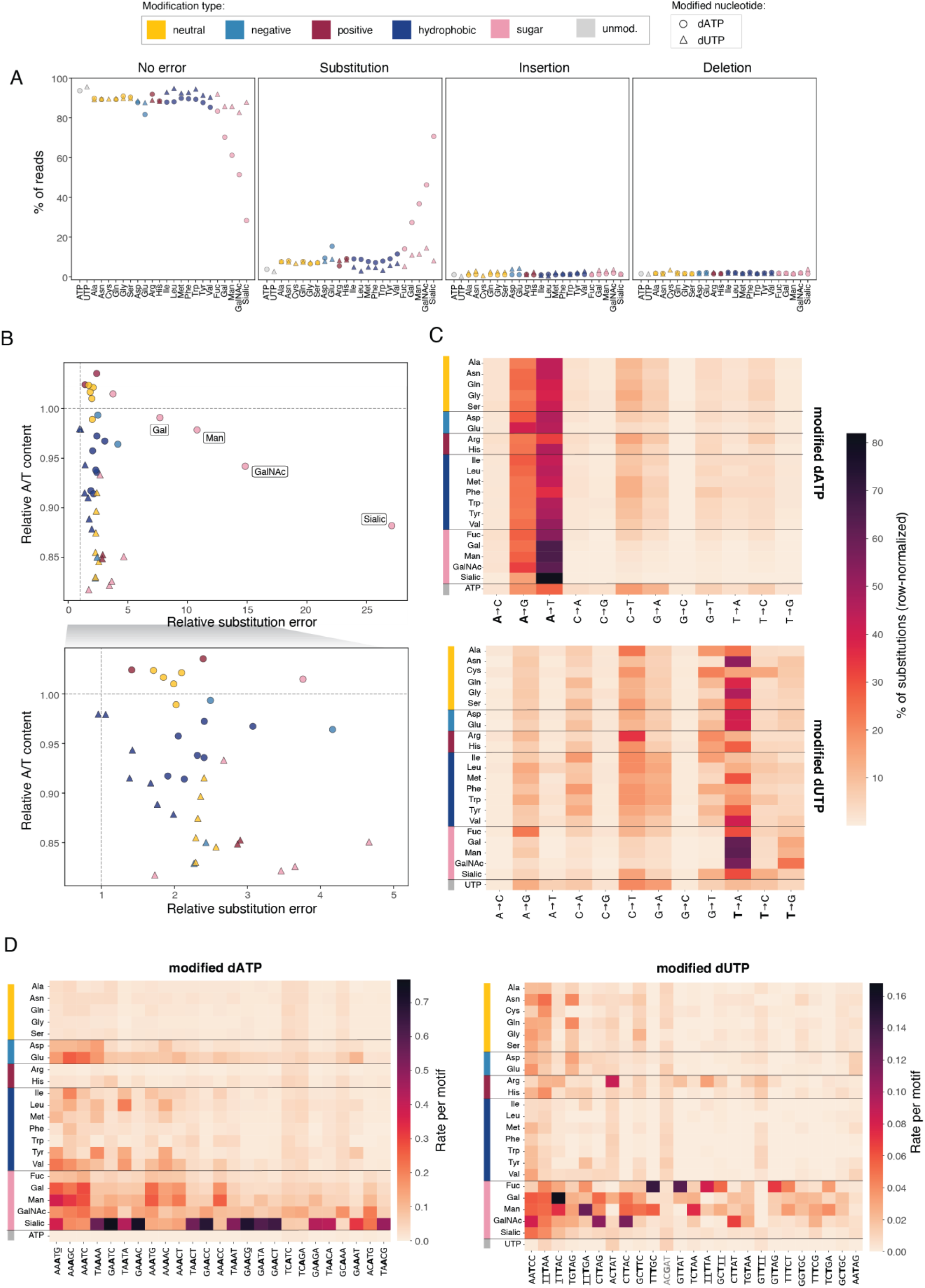
**A)** Fraction of assigned reads weighted by read abundance without errors, with at least one substitution, with at least one insertion, or deletion, as determined by Needleman-Wunsch alignment. Substitutions, insertions, and deletions are not mutually exclusive. Colors and shapes are as in Figure 1A, with unmodified nucleotides shown in grey. **B)** Relative, read weighted substitution rate per analyzed base considering only reads with substitutions and no insertions or deletions, plotted against relative adenine or thymine sequence content for assigned sequences from samples transcribed with modified dATPs or modified dUTPs, respectively. Both values are normalized to the corresponding unmodified controls. Bottom panel shows a zoomed view of substitution error rates between 0.5 and 5.2. **C)** Heatmap of base substitution pattern frequencies (expected base → sequenced base) for assigned reads of samples containing modified adenosines (top panel) and uridines (bottom panel). Only reads without deletion or insertion events were considered. For each modification or control (row), the frequency of each substitution pattern (read weighted) is calculated relative to the total number of substitution events for that sample. **D)** Heatmap showing the sequence context surrounding the observed substitutions (bold) detected in samples containing modified adenosines (left) or modified uridines (right), with unmodified ATP or UTP samples shown as control (bottom lanes). Substitutions were derived exclusively from reads containing no insertions or deletions. NNYNN motifs, where Y represents the substituted base, were filtered to retain motifs with more than 100,000 read-weighted substitution events across all samples. Motifs were ranked according to their mean relative substitution frequency across samples containing modified nucleotides, and the top 25 motifs are displayed. Flanking nucleotides that overlap with the constant regions adjacent to the 38 seed sequences (see Table S2) are underlined.

Across all samples, the decline in fidelity was primarily driven by substitutions (mean 8.5% of reads), while insertions (mean 1.4%) and deletions (mean 1.8%) were less frequent (Fig. 3A, S2A, B). For modified dUTPs, substitution only occurred in 9% of the analyzed reads, except for mannose, galactose, and GalNAc that was observed in 11–15% of the reads. Amino acid-modified dATPs showed similar substitution levels (5-15%), whereas glycosyl-modified dATPs (excluding fucose) exhibited substantially elevated substitution frequencies being detected in 27–70% of the reads.

Substitution rates were highly reproducible across independent replicates (Fig. S2B), confirming the robustness of our error profiling. For most modifications, substitution rates increased by less than five-fold relative to the controls, independently of the nucleobase (Fig. 3B). Amino acid-derived modifications consistently showed low error rates, rarely exceeding a 3-fold increase across both nucleobases, with Glu-dATP representing a minor outlier (4.2-fold increase). In contrast, sugar modifications on adenosines (galactose, mannose, GalNAc, and sialic acid) exhibited markedly elevated substitution rates, ranging from 8- to 27-fold above control levels (Fig. 3B).

These substitution error rates were directly reflected in the nucleotide composition of the respective sample pools. Generally, adenine content was largely preserved across most modified dATP samples relative to controls. However, glycosyl-modified dATPs generally (except Fuc-dATP) exhibited elevated substitution rates, leading to a reduction in adenine content of up to 12% (Fig. 3B, S2C). Negatively charged dATP modifications resulted in only minor decreases (up to 4%), while hydrophobic modifications caused reductions of 3-9 %, consistent with the higher divergence in sequence distribution previously observed for these specific modifications (Fig. S1H). Interestingly, positively charged and neutral amino acid modifications led to a slight increase in adenine content (up to 3%). In contrast, modified dUTPs exhibited a more pronounced depletion of thymine content across nearly all modifications, with reductions of up to 18% relative to the control. These widespread depletions mirror the broad sequence divergence observed in the library pool (Fig. S1H), where the most divergent sequence distributions corresponded to the strongest losses in thymine content and, consequently, losses of the modified bases.

We next analyzed substitution profiles to determine the nature of nucleotide misincorporations (Fig. 3C). Transversions of the modified base to the complementary base (A→T for modified dATPs and T→A for modified dUTPs) represented the predominant error types. These substitutions account for the observed depletion of adenine and thymine and the corresponding increase in their complementary bases in the sequenced pools (Fig. 3B & S2C). Interestingly, despite elevated A→G transitions, modified dATPs maintained stable guanine levels, whereas modified dUTPs were associated with an average 7% reduction in guanine content and a 12% increase in cytosine. As no dominant high-frequency substitution pattern directly explains this compositional shift (Fig. 3C), these changes likely arise from broader sequence divergence introduced during transcription and reverse transcription (Fig. S1H).

To determine whether substitutions occurred randomly or were influenced by specific sequence features, we analyzed the local +/-2 sequence positions (NNYNN) surrounding each substitution event. This analysis revealed that substitution hotspots are predominantly found within homopolymer stretches of at leat two consecutive modified nucleotides (Fig. 3D; dATP: 22 of top 25 motifs; dUTP: 15 of top 25). Notably, several motifs extended into the 5’-constant regions containing terminal thymines (e.g., denoted by underlined nucleotides in Fig. 3D, right panel). These homopolymer-associated errors were most pronounced for glycosyl-modified nucleotides.

### UMI-based analysis reveals RT as driver for substitutions

To determine whether the observed substitutions are introduced during transcription or reverse transcription, we applied a UMI-based deconvolution strategy adapted from Gout et al. (Fig. S3A, B) [54]. UMI-tagged transcripts of the highly represented sequence 19 (Fig. 2B, bottom panel) were generated using UTP, His-dUTP, Leu-dUTP, and Phe-dUTP during transcription. These modifications were selected due to their distinct chemical properties and observed differences in incorporation efficiency and error profiles. We performed three consecutive rounds of cDNA synthesis from the immobilized UMI-tagged transcripts, enabling us to distinguish between transcription-based errors (shared across multiple RT replicates) and errors introduced during downstream processing (RT, PCR or NGS). The analysis revealed that transcription-based errors were negligible, whereas most substitutions arose within individual RT rounds (Fig. S3C, D). Furthermore, sequencing of the DNA template showed a baseline substitution rate four- to six-fold lower than that observed for RNA samples (Fig. S3D), confirming that the substitutions observed in our workflow predominantly originate from errors introduced at the SSIII-mediated reverse transcription step rather than during IVT.

### In vitro selection of His-dUTP–modified aptamers

To test the applicability of base-modified nucleotides to sustain multiple rounds of *in vitro* selection and amplification, we subjected a fully His-modified-dUTP containing RNA library to Systematic Evolution of Ligands by EXponential enrichment (SELEX) to generate His-modified uridine containing aptamers.

To mitigate the amplification biases observed in our fidelity analysis, we utilized a revised library design featuring constant regions devoid of uridines. This ensures that the primer-binding sites remain unmodified, thereby maximizing IVT and RT recovery yields and simplifying potential downstream sequence truncation. As a target, we chose hemagglutinin (HA), a therapeutically relevant glycoprotein from the influenza H1 A/England/195/09 strain [55]. Six selection cycles were performed under progressively stringent conditions by adjusting incubation times, washing steps, and target concentrations (Fig. S4A). Counter-selection using empty beads and an unrelated His-tagged protein were performed to discard non-specific binders. After each cycle, bound sequences were recovered, reverse-transcribed, PCR-amplified, and used as input for the sub-sequent round. High-throughput sequencing of libraries from the initial pool and after cycles 2–6 revealed progressive enrichment of specific sequences (Fig. 4A, S4B), with the top 10 candidates reaching up to 40% enrichment after round 6 (Fig. 4B). Importantly, 6 rounds of iterative transcription-reverse transcription with His-dUTP did not result in any depletion of uridines in the library (Fig. S4C), indicating that even after multiple rounds of copying, His-dUTP incorporation does not impose a selection bias toward uridine-depleted variants.

**Figure 4:**
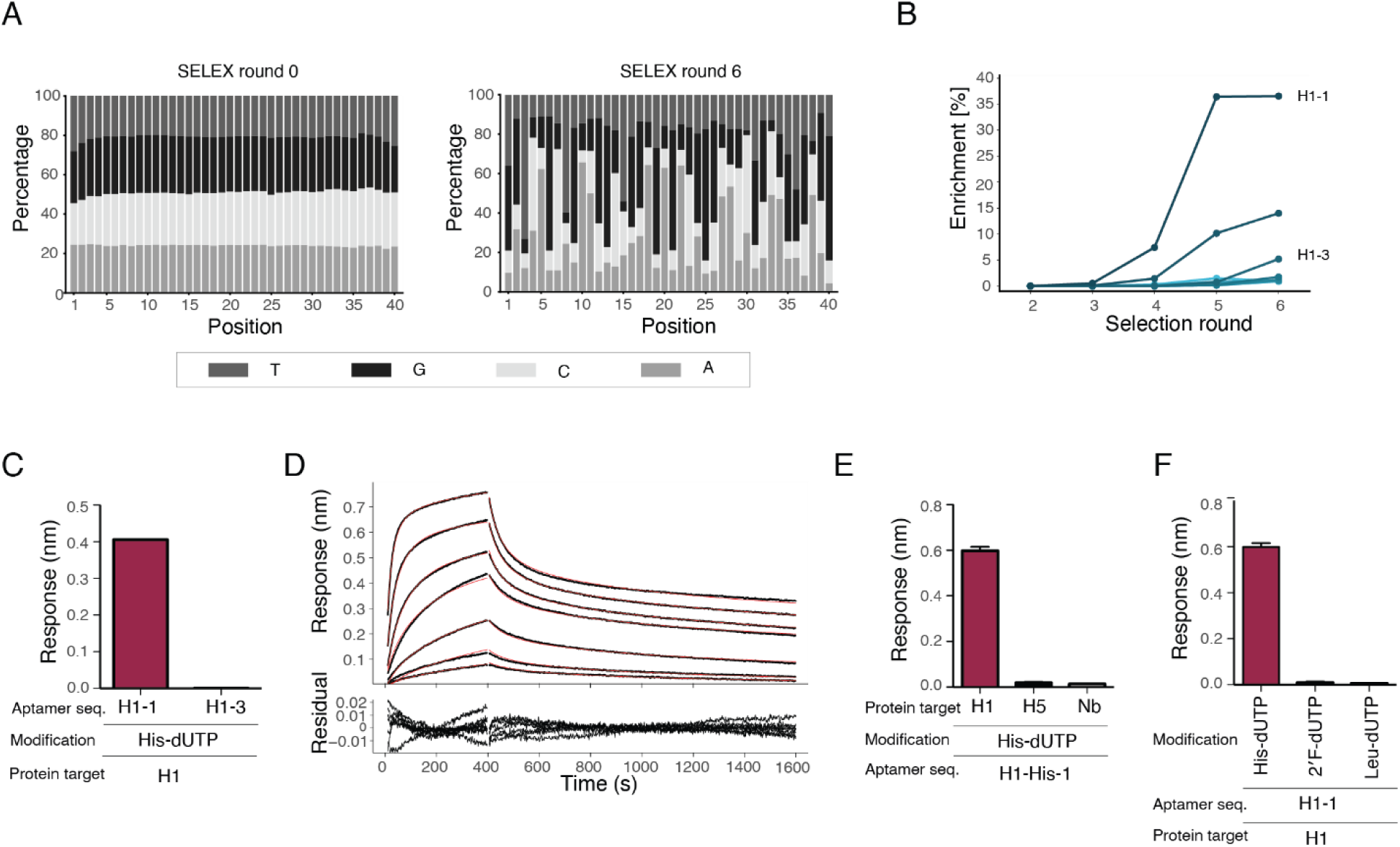
**A)** Nucleotide composition across the 40-nt random region of the starting SELEX library (round 0) and after selection round 6. Grey shades indicate nucleotide identities as shown in the legend. **B)** NGS profiling showing sequence enrichment (frequency, %) of the top 10 candidates across SELEX rounds 2–6. **C)** Screening of selected sequences for binding to H1 hemagglutinin using BLI. Average BLI response after 98 s of association for sequences H1-1 and H1-3 containing histidine-modified uridines, immobilized on streptavidin BLI sensors and exposed to H1 hemagglutinin in solution at 150 nM (n = 3). **D)** Binding kinetics of aptamer H1-His-1 to H1 hemagglutinin measured by BLI. The biotinylated aptamer was immobilized on streptavidin-coated BLI sensors and exposed to a two-fold dilution of H1 hemagglutinin in solution starting at 150 nM. Association phase: 0–400 s; dissociation phase: 400–1600 s. Global fitting of association and dissociation curves (red lines) was performed using Evilfit accounting for surface effects. Residuals are shown below. **E)** Assessment of H1-1 aptamer specificity. Average BLI response after 98 s of association for sequence H1-1 containing histidine-modified uridines, immobilized on streptavidin BLI sensors and exposed to 150 nM H1 hemagglutinin and H5 hemagglutinin or a 300 nM nanobody control in solution (n = 3). **F)** Evaluation of the impact of uridine modifications on H1-1 aptamer binding. Average BLI response after 98 s of association for H1-1 containing histidine-modified uridines or substituted with leucine- or 2F’-modified uridines, immobilized on streptavidin BLI sensors and exposed to 150 nM H1 hemagglutinin in solution (n = 3).

Following enrichment analysis and sequence motif clustering, two sequences (H1-His-1 and H1-His-3) were selected for binding characterization. Bio-layer interferometry (BLI) demonstrated that H1-His-1 bound specifically to the H1 trimer, whereas H1-His-3 showed no detectable interaction (Fig. 4C, S4D). H1-His-1 exhibited strong affinity in the low nanomolar range (K_D_ = 31.5 nM; Fig. 4D) and displayed high specificity, showing no binding to a related HA protein (H5 A/whooper swan/Mongolia/244/2005) or to the His-tagged nanobody used for counter-selection during SELEX (Fig. 4E, S4E). To assess the functional contribution of the histidine modification, H1-His-1 was transcribed with alternative modifications (2′F-dUTP or Leu-dUTP). None of these variants bound to H1 (Fig. 4F, S4F), demonstrating that the histidine modification is essential for aptamer-target recognition.

## DISCUSSION

Base modifications constitute a key strategy for expanding the functional potential of RNA by conveying new properties such as improved stability, target binding affinity or more complex catalytic centers. In this study, we provide a systematic evaluation of Y639F mutant T7 RNA polymerase-mediated co-transcriptional incorporation of 23 chemically diverse base modifications—including amino acids and sugars differing in charge, polarity, hydrophobicity, and steric bulk—introduced at two distinct nucleobase positions on deoxynucleotides: the C5 position of dUTP and the C7 position of dATP. In addition, 2’F-dCTP was systematically used in all RNAs to test the compatibility and synergistic effect of this ribose modification, crucial for enhanced nuclease resistance, in combination with the reported base and deoxyribose modifications in our study. Importantly, in contrast to published work investigating the ability of T7 RNA polymerase to incorporate C5-substituted pyrimidines and 7-deazapurine analogs, we include detailed assessment of fidelity throughout transcription and reverse transcription [39]. This information is fundamental for workflows requiring robust and/or iterative amplification/copying such as in in vitro evolution.

Our data demonstrate that the Y639F mutant of T7 RNA polymerase efficiently incorporates the majority of base–modified nucleotides carrying amino acid and sugar functional groups. Although transcription yields declined substantially for certain combinations of nucleobase and modification, a notable anti-correlation between adenosine and uridine modification efficiency was observed. For instance, the clinically relevant targeting moiety GalNAc was incorporated with near-quantitative efficiency when conjugated to adenosine, but with less than 10% efficiency when attached to uridine. Conversely, hydrophobic amino acids such as leucine, isoleucine, tryptophan, and phenylalanine were incorporated far more efficiently when conjugated to uridine than to adenosine. These findings highlight that modification performance cannot be generalized across nucleobases and must be empirically assessed. Consistent with other reports [20], we also observe reduced efficiency for bulkier modifications, especially when using adenosines, underscoring a recurring limitation of polymerase promiscuity and substrate discrimination.

A central concern when transcribing RNA with non-canonical nucleotides is the potential introduction of sequence errors, including substitutions, insertions, and deletions. To address this, we analyzed sequence accuracy for a diverse pool of sequences following one round of transcription-reverse transcription using next-generation sequencing. Overall, high sequence accuracy was observed for modified nucleobases, although sugar-like modifications exhibited higher error rates for both adenosines and uridines. Since this strategy only reports on the combined transcription-reverse transcription fidelity, we further implemented a UMI-based deconvolution approach to disentangle these two processes. Focusing on representative amino acid–modified uridines (His, Leu, and Phe), we found that the majority of observed errors originated during the reverse transcription step rather than during transcription by T7 RNA polymerase. This strongly indicates that the T7 RNA polymerase incorporates the modified nucleotides with very high accuracy. Whether this conclusion extends to all modifications tested remains an important question for future studies.

Although overall transcription-reverse transcription fidelity was high for most modifications, notable exceptions were observed, particularly for sugar-modified adenosines. Substitution rates of up to ∼3% were detected for sialic acid–modified adenosine, likely reflecting the combined effects of steric bulk and negative charge associated with this branched sugar. One plausible explanation for the generally inferior performance of sugar modifications compared to amino acid modifications is their mode of attachment: Whereas amino acids are linked through a flexible C4 spacer, sugar modifications are directly conjugated through a shorter C2 spacer to the nucleobase, potentially increasing steric clashes with the active site of either the polymerase or the reverse transcriptase.

Based on our findings, several practical guidelines emerge. First, a high density of modifications near the 5′ end of transcripts should be avoided [44], as polymerase stalling or premature dissociation during transcription initiation might increase the formation of abortive products. Second, for sugar modifications, C5-modified uridines are generally preferred due to higher yields and lower error rates. Third, homopolymer stretches should be avoided or minimized, as they represent hotspots for sequence distortions and substitution errors. Finally, wherever possible, sugar-modified adenosines should be avoided due to their consistently lower fidelity.

The ability to both accurately transcribe and reverse transcribe highly modified RNA molecules is particularly important for in vitro selection workflows. While base-modified nucleic acids have long been explored in DNA aptamer selection, for example SOMAmers using amide-linked side chains [22,56], libraries incorporating click-modified bases for post-transcriptional functionalization [57], analogous strategies for RNA remain comparatively underdeveloped. The use of RNA offers additional advantages, including enhanced nuclease resistance compared to DNA when using 2’-fluoro and 2’-deoxy modifications, and potentially increased structural diversity and robustness via hydrogen-bonding potential through 2′-OH groups at the remaining [58]. To demonstrate the applicability of our system, we selected histidine-modified uridine aptamers against influenza hemagglutinin. Histidine is particularly attractive due to its pKa near physiological pH, allowing protonation and modulation of charge at mildly acidic conditions, thereby enabling potential pH-sensitive binding. Over six rounds of selection, only limited loss of uridine content was observed, culminating in the identification of a highly specific aptamer with low-nanomolar affinity. This demonstrates that chemically modified RNA libraries generated by our method are fully compatible with in vitro evolution.

In summary, our study provides a comprehensive framework for enzymatic synthesis of chemically diverse RNAs with high efficiency and accuracy and defines practical design principles for their implementation. In future work, systematic investigation of how the reported modifications affect enzymatic stability, as well as the biophysical and thermodynamic properties of the resulting oligonucleotides, will further refine these design principles and enable the discovery of new RNA functions. While the showcased aptamer selection highlights one potential application towards the in vitro selection of RNAs with expanded chemical repertoire, the presented approach can for example be combined with rationally designed RNA nanostructures to display functional groups at predefined positions, thereby engineering optimal interaction surfaces for multivalent or heterovalent binding events often required for cooperative recognition of proteins or cell-surface receptors [59]. Exploring this approach to the development of chemically modified therapeutic RNAs, either through high-throughput library generation or by enabling modification of long RNAs (e.g., mRNAs) beyond the reach of current solid-phase synthesis methods, could pave the way for new avenues in RNA engineering and therapeutics. Together, these features underscore the broad potential of this platform for applications in nanotechnology, synthetic biology, and RNA-based therapeutics.

## DATA AVAILABILITY

Raw sequencing and mass analysis data is available from the authors upon reasonable request.

## SUPPLEMENTARY DATA

Supplementary Data are available at NAR online.

## AUTHOR CONTRIBUTIONS

Julian Valero: Conceptualization, Formal analysis, Methodology, Validation, Writing—original draft, review & editing. Kevin Neis: Methodology, Data analysis, Writing—original draft. Laia Civit: Methodology, Data analysis, Writing. Søren Fjelstrup: Data analysis. Maria Gockert: Writing—review & editing. Jørgen Kjems: Conceptualization, fund raising, writing—review & editing.

