## Supplementary Information for "Expanding the chemical diversity of RNA by transcriptional incorporation of amino acid- and glycosyl-modified nucleotides"

### Supplementary Figures

#### Supplementary Figure S1:

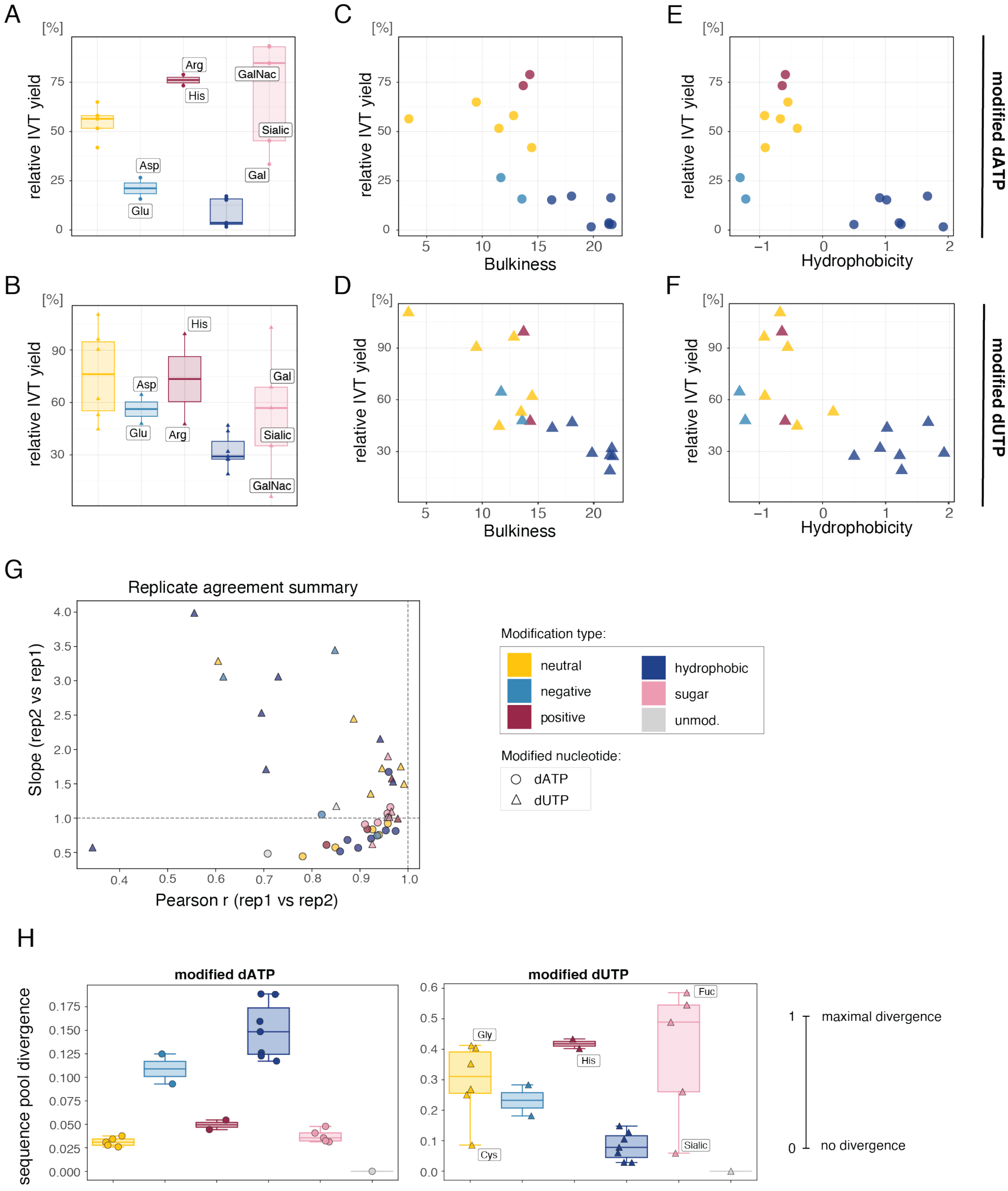

A) Relative IVT yields from the pooled DNA templates using modified dATPs, grouped by modification type. For all panels, color code represents modification type, with neutral amino acid modifications in yellow, negatively charged in light-blue, positively charged in dark-red and hydrophobic in dark-blue. Sugar-like modifications are pink. Modified adenosins are indicated by round points, modified uridines by triangles.

B) Relative IVT yields from the pooled DNA templates using modified dUTPs, grouped by modification type.

C) Relative IVT yield using amino acid-modified dATPs as a function of amino acid bulkiness.

D) Relative IVT yield using amino acid-modified dUTPs as a function of amino acid bulkiness.

E) Relative IVT yield using amino acid-modified dATPs as a function of amino acid hydrophobicity.

F) Relative IVT yield using amino acid-modified dUTPs as a function of amino acid hydrophobicity.

G) Pearson correlation coefficient (r) plotted against the slope of a linear fit comparing relative seed abundance profiles of assigned sequencing reads from duplicate experiments. Values closer to 1 indicate higher agreement between replicates.

H) Jensen–Shannon divergence (JSD) showing the difference in sequence composition for samples with modified adenosines (left panel) and modified uridines (right panel) compared to respective unmodified controls (ATP, UTP). Only sequences within a Levenshtein-Damerau distance of ≤5 from one of the 38 respective seed sequences were considered (from now termed as assigned sequences). A JSD value of 0 indicates identical sequence distribution, while a value of 1 indicates maximal divergence.

**Supplementary Figure S2:**

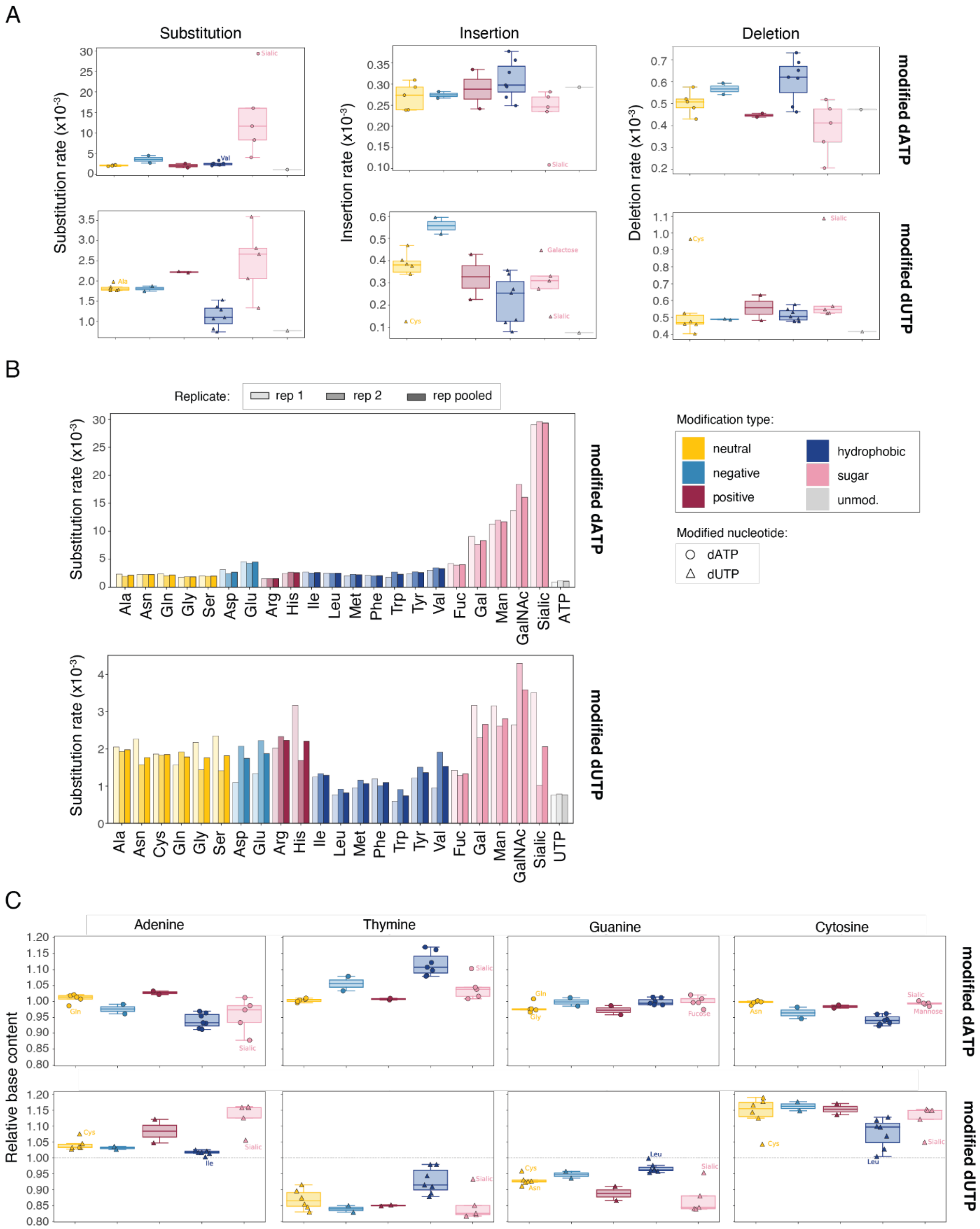

A) Read weighted error rates per analyzed base for subsitiution (left), insertion (mid), and deletion (right) events in assigned sequences containing exclusive error types, grouped by modification type for samples containing modified adenosiones (top) or uridines (bottom), as well as unmodified ATP and UTP controls, respectively. Boxes indicate the interquartile range (IQR) with the median shown as a line, whiskers extend to most extrem values within 1.5x IQR, and outliers are indiviually lablled with sample names. Colors and shapes are as in Figure 2A.

B) Exclusive, read weighted substitution error rates in assigned reads for samples containing modified or unmodified adenosines (top panel) and uridines (bottom panel), calculated for individual replicates (rep1, rep2) and after pooling replicates (rep pooled).

C) Relative nucleotide content of assigned sequences from samples transcribed with modified dATPs(top) or dUTPs (bottom), relative to the respective unmodified contol samples (ATP or UTP). Plotting as in Figure S2A.

**Supplementary Figure S3:**

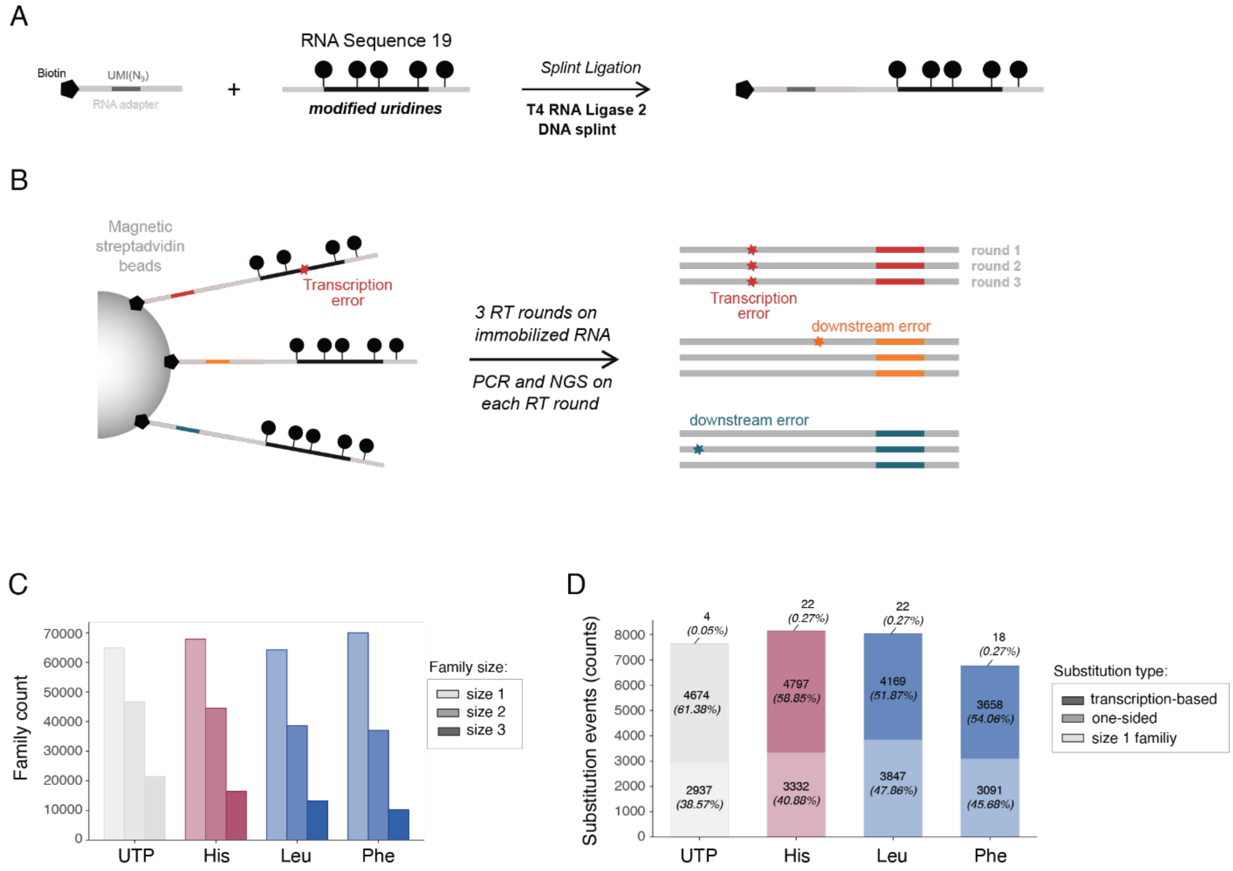

A) Schematic workflow for generating cycling reverse transcription templates. Using T4 RNA ligase and a DNA splint, an exemplary RNA sequence from the pool of 38 sequences (sequence 19) containing modified uridines (circles) was ligated to a 5' biotin-labeled RNA adapter containing a 9-nucleotide unique molecular identifier (UMI).

B) Schematic overview of cycling reverse transcription and error classification. Biotinylated templates generated as in A) are immobilized on magnetic streptavidin beads and subjected to reverse transcription (RT), PCR amplification, and NGS for three consecutive cycles. UMI-guided grouping of NGS reads identifies sequence groups derived from the same ancestor molecule. For groups with two or three members, errors (stars) shared by at least two members of a group are classified as IVT-derived transcription errors, whereas errors found in only a single member are classified as downstream, non–IVT-related errors.

C) Number of sequence families (defined by UMI) detected in only one cycling round (size 1), two rounds (size 2), or all three rounds (size 3). RNA template sequence 19 was generated by IVT using either UTP or dUTPs modified with histidine (His), leucine (Leu), or phenylalanine (Phe). Colors indicate modification class, with red representing positively charged and blue hydrophobic modifications.

D) Total number and frequency of substitutions in sequences with no indels relative to template sequence 19 in samples containing unmodified uridines or histidine-, leucine-, or phenylalanine-modified uridines. Substitutions are classified according to their representation within sequence families: size 1 family substitutions occur in families detected in only one cycling round; one-sided substitutions occur in a single sequence within a size 2 or size 3 family; and transcription-based substitutions are detected in at least two sequences within a size 2 or size 3 family.

**Supplementary Figure S4:**

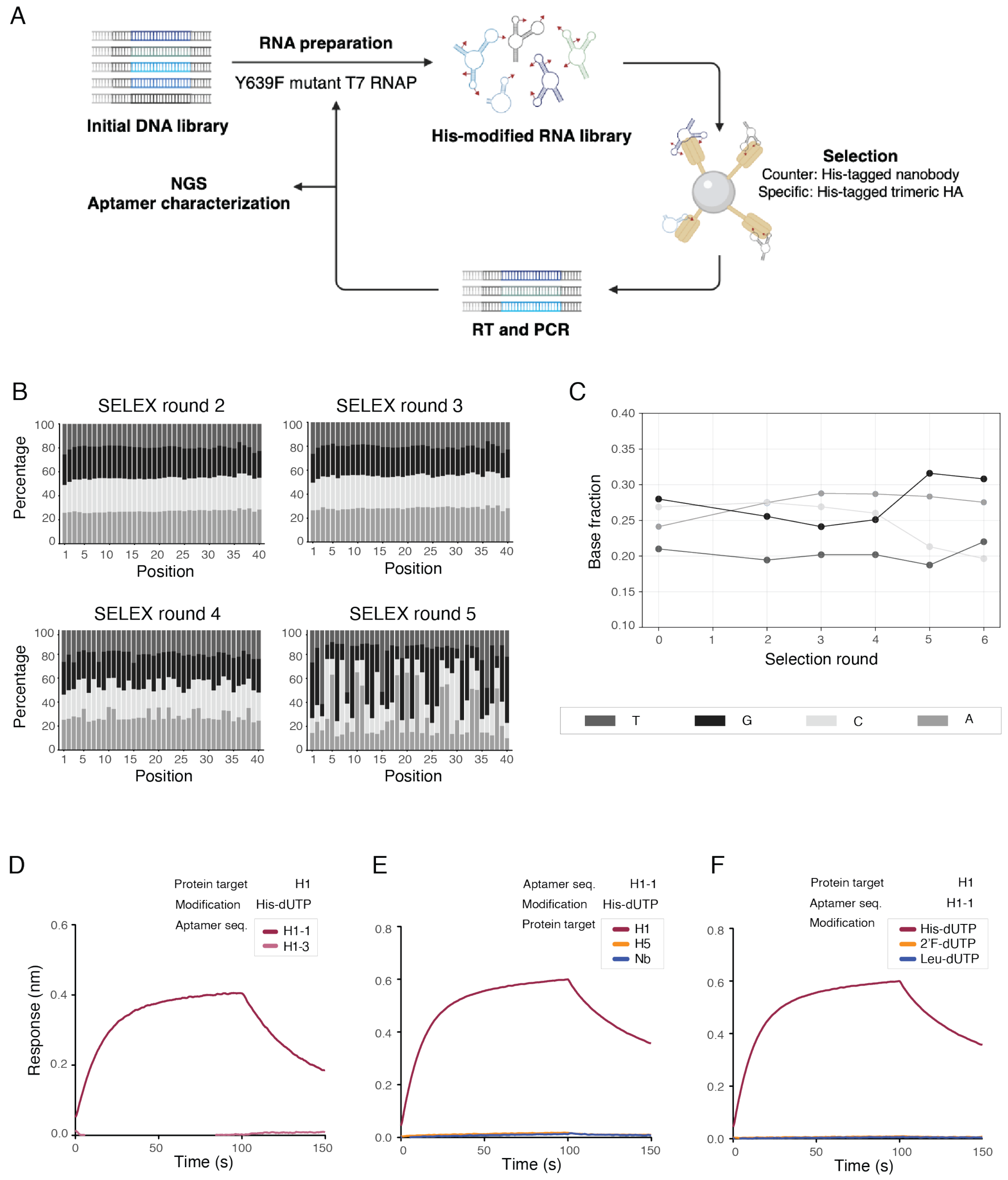

A) SELEX scheme for the selection of His-modified aptamers against H1 hemagglutinin.

B) Nucleotide composition across the 40-nt random region of selection rounds 2-5. Grey shades indicate nucleotide identities as shown in the legend.

C) Overall nucleotide compositions across all sequences within the SELEX rounds. Grey shades indicate nucleotide identities as shown in the legend.

D-F) Sensograms of the BLI measurements shown in Figure 4, with the left panel (D) corresponding to the data displayed in Figure 4C, the middle panel (E) to Figure 4E, and the right panel (F) to Figure 4F.

#### Supplementary Tables

**Table S1.** Mass analysis (theoretical and experimentally found m/z values) of modified dATP and dUTP nucleotides.

| | dATP mass | | dUTP mass | |
|---|---|---|---|---|
| | **Theoretical** | **Exper. found** | **Theoretical** | **Exper. Found** |
| Leu | 755.14 | 755.15 | 733.10 | 733.12 |
| Met | 773.09 | 773.10 | 751.06 | 751.07 |
| Trp | 828.13 | 828.14 | 806.11 | 806.11 |
| Tyr | 805.12 | 805.13 | 783.08 | 783.10 |
| Phe | 789.12 | 789.13 | 767.09 | 767.10 |
| Val | 741.12 | 741.13 | 719.09 | 719.10 |
| Ile | 755.14 | 755.15 | 733.10 | 733.12 |
| Asp | 757.08 | 757.09 | 735.05 | 735.06 |
| Glu | 771.10 | 771.11 | 749.06 | 749.08 |
| Ser | 729.08 | 729.10 | 707.05 | 707.06 |
| Gln | 770.11 | 770.12 | 748.08 | 748.09 |
| Ala | 713.09 | 713.10 | 691.06 | 691.07 |
| Asn | 756.10 | 756.11 | 734.06 | 734.08 |
| Gly | 699.07 | 699.08 | 677.04 | 677.05 |
| Thr | 743.10 | 743.11 | 721.07 | 721.08 |
| Arg | 798.15 | 798.16 | 776.12 | 776.13 |
| His | 779.11 | 779.12 | 757.08 | 757.09 |
| Sialic acid | 847.12 | 847.12 | 825.09 | 825.09 |
| Mannose | 762.10 | 762.11 | 740.07 | 740.07 |
| GalNac | 803.13 | 803.13 | 781.10 | 781.10 |

| | | | | |
|---|---|---|---|---|
| Galactose | 762.10 | 762.10 | 740.07 | 740.07 |
| Fucose | 746.11 | 746.11 | 724.08 | 724.08 |

**Table S2.** Appendix (sequences)

Constant region: 5′- GGGGCCACCAACGACATT-Nº seq-GTTGATATAAATAGTGCCCATGGATCC

| | |
|---|---|
| Seq1 | GGTTTCAATGTCCCGGTACTGTCATCACATACAATG |
| Seq2 | GCTTATCGTGTCTTGGTACTGTCAGCACATACTATG |
| Seq3 | GTCTAACTGTCGGAGGTACTGTCAACACATACGATG |
| Seq4 | GCGGTGCTTCGCTGCGACATCCGCGAAACCAACGTC |
| Seq5 | TGAGTGGTGCTCCTACAACGCAGGAACCTCAATGCT |
| Seq6 | CCGACCTTAGTCAATCACTTCGTTCGATGAATAGCA |
| Seq7 | AGTGTAGGAACTTTCGGACCTCCATCATAAGCATGC |
| Seq8 | AGTACACCGTAACACTACGGCTTATCATAAGCATGAC |
| Seq9 | GCATCACCGCTCGACTTACATCGACCCGTGTAAAAC |
| Seq10 | ACTGGATGTTTGCTTAGTCCAGTGATAGCACATGC |
| Seq11 | AGCAACTACCATGACAACGTGTTAGTCGGTCGAAAT |
| Seq12 | GAACCACGTAGGCTCGTTTCTGAGCCGATCTCGAT |
| Seq13 | TGCGACTGTTATAACCTAACAGCGACGTAAAGATA |
| Seq14 | TAACGAATATGATCCGATTCGCTCTCAGCACATCAC |
| Seq15 | AACGTTAAATGAGCCAAGCGACTTGCATACTTGGAC |
| Seq16 | TATCGAATTGATAACCTTACGCGAGAGCGTAGTTC |
| Seq17 | AGTGTGTACGAACAACGTTCGTCCCATCATAAGCAT |
| Seq18 | TTACCTACTCCGTACATTCGGATGGTGCCGAACGTT |
| Seq19 | CGAGCGGTAAATCCAAGCCGCACTTCAGTCCGTCAC |
| Seq20 | GTTCGATCGGCGTTATAATACCACGCGACCCTTGAC |
| Seq21 | TAAAGCGTCGGGCCGTACTGTCAATACATACAATG |
| Seq22 | ACGCGTCTTGGTGACGCTTTTGTGAATCTAAGCCAC |
| Seq23 | TGGTCGCGGGGATTGCGCAAATGTTGTAACTGAAC |
| Seq24 | TTTTAGCTCTGATCTAAGCTAAACTCACGCCATCCC |
| Seq25 | TTGCGACGATTATTGAAACCACACGTCGCCTCAGGT |
| Seq26 | CAAAGCCGACATGCTGGTGTTCATGAATACACCATC |
| Seq27 | AAATTTCCCTAAATTGTAGGCAGGTACACCGTACCC |
| Seq28 | TTACGGTACGGCTAAAACTGCACGACCTCTCAGGCC |
| Seq29 | TTAGGTGCTCTGAACTTTACATCGGAACTAACGTTG |
| Seq30 | CTTAACATCAGCACAAATTCTAGTGCGCATTCTGCC |
| Seq31 | TTTTCCCTTCTCAGTAAGGCAGTCGTGTTTCATGACC |
| Seq32 | TCGTTCGATCTCTCGAGTTCTCAGAGCGACCTTTAT |
| Seq33 | TCGGCGTTGTCAGAAGACCGAAACTTAGACTGTGTT |
| Seq34 | TGAGATACCAGGCGGTCATATGACACCGAACTAGTC |
| Seq35 | TTCGCGCAGAGCTTCAGGTTTCAATTTAACCCTTTC |
| Seq36 | CTTGTCATCAGAAACGCGCTAGTTTCGCATTCTGTC |

Seq37 TTTAGAAGTTGATCTAATTCTAACTCAGTACGTCAC
Seq38 TTTCCGCTGAGTAAAGCGCAGTCGAGTCTTCCGACC

**Table S3.** Sequences synthesized by Integrated DNA Technologies (IDT).

| name | sequence | length |
|---|---|---|
| KKfw | CGCGGATCCTAATACGACTCACTATAGGGGCCACCAACGACATT | 44 |
| KKrv | CCCGACACCCGCGGATCCATGGGCACTATTTATATCAA | 38 |
| KKrv_short | GATCCATGGGCACTATTTATATCAAC | 26 |
| KKfw_adapter | GTGTATAAGAGACAGATCC | 19 |
| KKfw_extension | TCGTCGGCAGCGTCAGATGTGTATAAGAGACAGATCC | 37 |
| KKrv_extension | GTCTCGTGGGCTCGGAGATGTGTATAAGAGACAGGATCCATGGGCACTATTTATATCAAC | 60 |
| KK_RNA_BioAdapt | /5Biosg/rGrUrGrUrArUrArArGrArGrArCrArGrArUrCrCrNrNrNrNrNrNrNrNrNrGrCrCrGrArGrUrGrCrArGrC | 38 |
| DNA splint | GTCGTTGGTGGCCCCGCTGCACTCGGC | 27 |

**Table S4.** Sequence analysis of most and least abundant seed sequences.

| Base | Category | # of samples | Seed ID | Sequence | A count | max A run |
|---|---|---|---|---|---|---|
| ATP | More abundant | 19 | 34 | TGAGATACCAGGCGGTCATATGACACCGAACTAGTC | 11 | 2 |
| | | 10 | 14 | TAACGAATATGATCCGATTCGCTCTCAGCACATCAC | 11 | 2 |
| | | 9 | 29 | TTAGGTGCTCTGAACTTTACATCGGAACTAACGTTG | 9 | 2 |
| | | 7 | 12 | GAACCACGTAGGCTCGTTTCTGAGCCGATCTCGAT | 7 | 2 |
| | Less abundant | 19 | 13 | TGCGACTGTTATAACCTAACAGCGACGTAAAGATA | 13 | 3 |
| | | 8 | 17 | AGTGTGTACGAACAACGTTCGTCCCATCATAAGCAT | 11 | 2 |
| | | 6 | 4 | GCGGTGCTTCGCTGCGACATCCGCGAAACCAACGTC | 7 | 3 |
| | | 6 | 11 | AGCAACTACCATGACAACGTGTTAGTCGGTCGAAAT | 12 | 3 |
| | | 6 | 35 | TTCGCGCAGAGCTTCAGGTTTCAATTTAACCCTTTC | 7 | 2 |
| Base | Category | # of samples | Seed ID | Sequence | T count | max T run |
| UTP | More abundant | 19 | 19 | CGAGCGGTAAATCCAAGCCGCACTTCAGTCCGTCAC | 6 | 2 |
| | | 16 | 6 | CCGACCTTAGTCAATCACTTCGTTCGATGAATAGCA | 10 | 2 |
| | | 7 | 7 | AGTGTAGGAACTTTCGGACCTCCATCATAAGCATGC | 9 | 3 |

| | | | | | | |
|---|---|---|---|---|---|---|
| | | 7 | 8 | AGTACACCGTAACACTACGGCTTATCATAAGCATGAC | 8 | 2 |
| | | 7 | 17 | AGTGTGTACGAACAACGTTCGTCCCATCATAAGCAT | 9 | 2 |
| | Less abundant | 15 | 13 | TGCGACTGTTATAACCTAACAGCGACGTAAAGATA | 8 | 2 |
| | | 13 | 35 | TTCGCGCAGAGCTTCAGGTTTCAATTTAACCCTTTC | 13 | 3 |
| | | 11 | 32 | TCGTTCGATCTCTCGAGTTCTCAGAGCGACCTTTAT | 13 | 3 |

*Not found only Arg, Fuc, Gal (Sequence 35 on As). Included in 6 appearances

**Table S5.** ssDNA for SELEX experiment

| name | sequence | length |
|---|---|---|
| N40_mU_ssDNA | TCGGGCGTGTCGTCTG-N40-CCGCTTCCTCCTCCC | 71 |
| N40_mU_fw | TAATACGACTCACTATAGGGAGGAGGAAGCGG | 32 |
| N40_mU_rv | TCGGGCGTGTCGTCTG | 16 |
| N40-mU NGS "overhang fw" | TCGTCGGCAGCGTCAGATGTGTATAAGAGACAGTAATACGACTCACTATAGGGAGGAGGAAGCGG | 65 |
| N40-mU NGS "overhang rv" | GTCTCGTGGGCTCGGAGATGTGTATAAGAGACAGTCGGGCGTGTCGTCTG | 50 |
| Hi_His_1 | TCGGGCGTGTCGTCTGCCTCACTTCCGGTTCCCGTCTCTCCAGCCTTCAGCTGACCCCGCTTCCTCCTCCC | 71 |
| Hi_His_3 | TCGGGCGTGTCGTCTGCGGATTAGTATTACCCAACACCCGATATCCCAACCCTATACCGCTTCCTCCTCCC | 71 |
